# Pre-mRNA processing entropy in a mouse model of trauma

**DOI:** 10.1101/2021.01.25.428080

**Authors:** Maximilian S. Jentzsch, Alger M. Fredericks, Jason T. Machan, Alfred Ayala, Sean F. Monaghan

## Abstract

**Purpose:** Next generation sequencing has expanded our understanding of many disease processes, including trauma and critical illness. Many studies focus identifying a small set of genes or proteins that are aberrantly expressed. Our objective was to determine whether global differences in pre-mRNA processing entropy, or disorder, could offer novel insights in the setting of critical illness.

**Methods:** We used an established murine model of trauma that consisted of hemorrhagic shock and cecal ligation and puncture. In our first experiment mice exposed to trauma were compared to controls. In our second experiment, survival 14 days after exposure to trauma was studied. Using deep RNA sequencing we determined entropy values for every pre-mRNA processing event identified. We then used principal component analysis (PCA) to conduct unsupervised classification of the data.

**Results:** Mice exposed to trauma separated from controls using PCA. Similarly, mice that did not survive 14 days post exposure clustered closely together on PCA.

**Conclusion:** Our results suggest that there is a substantial difference in global pre-mRNA processing entropy in mice exposed to trauma vs. controls, and that pre-mRNA processing entropy may be helpful in predicting mortality. The method introduced here is easily transferrable to other disease processes and samples.

## Introduction

Trauma is the leading cause of death up to age 46 in the United States.^1^ Death from trauma typically occurs in one of three scenarios: immediately on scene or in the pre-hospital setting, within 24 hours, or within days to weeks of the inciting event(s)^2^. Outcomes vary widely among patients who survive 24 hours post injury. Patients with comparable injuries, comorbidities, and laboratory values may have drastically different outcomes with no clear physiological explanation. Early identification of markers of survival or disease severity could improve outcomes on a multitude of levels of intervention strategies, from triage to early initiation of personalized treatment and beyond.

The development of next generation sequencing has offered novel insight into the pathogenicity of critical illness, including trauma. Many studies on this topic have focused on identifying differences in gene expression.^3^ Less is known about other mechanisms that influence protein expression, such as alternative splicing.

In eukaryotic cells, pre-mRNA undergoes processing events occurring both co-transcriptionally and post transcriptionally. Alternative splicing is the process in which introns are removed and exons are ligated, yielding mature mRNA that is then ultimately translated into protein. Pre-mRNA splicing is a tightly regulated process that is spatially and temporally regulated. Understanding the potential effects of trauma on alternative pre-mRNA processing events may offer key insight into the diagnosis and treatment of trauma patients. Previous studies have reported that fever, hypoxia, and osmotic stress can influence pre-mRNA processing ^4–6^. Recent work from our group in an established animal model of trauma / Acute Respiratory Distress Syndrome (ARDS) has demonstrated that alternative pre-mRNA processing influences protein isoform distribution in critical illness ^7–9^.

Here, we offer a novel approach to the analysis of the effects of trauma on the basic molecular process of pre-mRNA processing. Specifically, we use entropy, the level of uncertainty in a random variable, to evaluate changes in pre-mRNA processing events. We demonstrate the application of this method for predicting trauma exposure and survival in a mouse model of trauma.

## Methods

Two approaches were used in this study. First, we evaluated differences in pre-mRNA processing entropy in mice 24 hours after the induction of trauma via hemorrhagic shock followed by cecal ligation and puncture (Hem/CLP) versus a sham model (detailed in methods section below). Second, we assessed differences in alternative pre-mRNA processing in mic at the time of trauma of mice who survived 14 days after exposure to Hem/CLP versus those who did not survive. The first approach consisted of three mice exposed to trauma and three controls; the second approach consisted of ten mice, all exposed to trauma.

### Mouse Model of Trauma

C57BL/6J mice from The Jackson Laboratory (Bar Harbor, ME) were used. Experiments were performed in accordance with National Institutes of Health guidelines, approval from the Animal Use Committee of Rhode Island Hospital (Providence, RI; AWC# 0079-13) and with consideration to the ARRIVE guidelines developed by the National Center for the Replacement, Refinement, and Reduction of Animals in Research ^10^. Severe trauma was induced by hemorrhagic shock followed by cecal ligation and puncture (CLP). This method has been previously demonstrated by our group ^8,11–13^. Briefly, mice were anesthetized in the supine position. Catheters were inserted in both femoral arteries and the mice were bled over a five to ten-minute period to a mean blood pressure of 30 mmHg (± 5mmHg). To achieve this level of hypotension, the mice typically had 1 mL of blood withdrawn, or approximately 50% of their total blood volume. This translates to a class 4 hemorrhagic shock in humans. The mice were kept stable for 90 minutes and then resuscitated intravenously with Ringers lactate at four times the drawn blood volume. For the control mice, sham hemorrhage was performed by ligating the femoral arteries without the subsequent blood draw, mimicking tissue destruction. On the following day, sepsis was induced in the mice by cecal ligation and puncture, which mimics a missed bowel injury on assessment of a human trauma patient.

For the second approach (the survival experiment), survival was determined at 14 days post CLP.

### Blood Samples and Sequencing

Blood samples were drawn at 24 hours after CLP for the exposure experiment and at the time of hemorrhage in the survival experiment. RNA from the blood samples was extracted using the MasterPure Complete RNA Purification kit for mice (epicenter, Madison, WI). The high concentration of globin RNA in blood samples required further processing with the GLOBINclear Kit (epicenter, Madison, WI). For deep sequencing, at least 1400 nanograms of RNA were required and all samples passed quality control standards. Samples were sequenced at Genewiz with a goal of at 40-80 million reads per sample.

### Alignment and Entropy calculations

RNA was sent to Genewiz, Inc. for deep sequencing. Sequencing results were subsequently run through FastQC as a quality control measure. The sequencing data was then aligned using STAR aligner with GRCm38/mm10 annotation downloaded from the UCSC genome browser. After confirming the quality of the raw data and initial alignment, the sequencing files were analyzed using Whippet.^14^ Whippet output contains several sub-routines that quantitate the levels of changes in pre-mRNA processing within RNA sequencing data, including entropy measures for all pre-mRNA processing events detected throughout the transcriptome. These measures were then compared between the murine model of trauma and sham control group in a principal component analysis.

### Principal Component Analysis

The output from Whippet yielded up to 221,010 raw entropy values per sample. A principal component analysis (PCA) was conducted to reduce the dimensionality of the dataset and to obtain an unsupervised classification of trends in entropy values among the samples. Raw entropy values from all samples were concatenated into one matrix and missing values were replaced with column means. The PCA was conducted using the *prcomp* function in base R, and the first two principal components were plotted against each other using *ggplot2* in R^15,16^. The percent variability explained was also calculated using *prcomp*. Outcome identifiers were added to the plots after the principal components were calculated; they were not part of the PCA. We conducted two PCAs to compare mice exposed to trauma vs. controls, and to compare mice that survived vs. died within 14 days after trauma.

## Results

Three mice exposed to the trauma model were compared to three mice in the control group (total n = 6). When plotting the first two principal components against each other, the exposed mice separated from the control mice (Figure 1).

**Figure 1.**
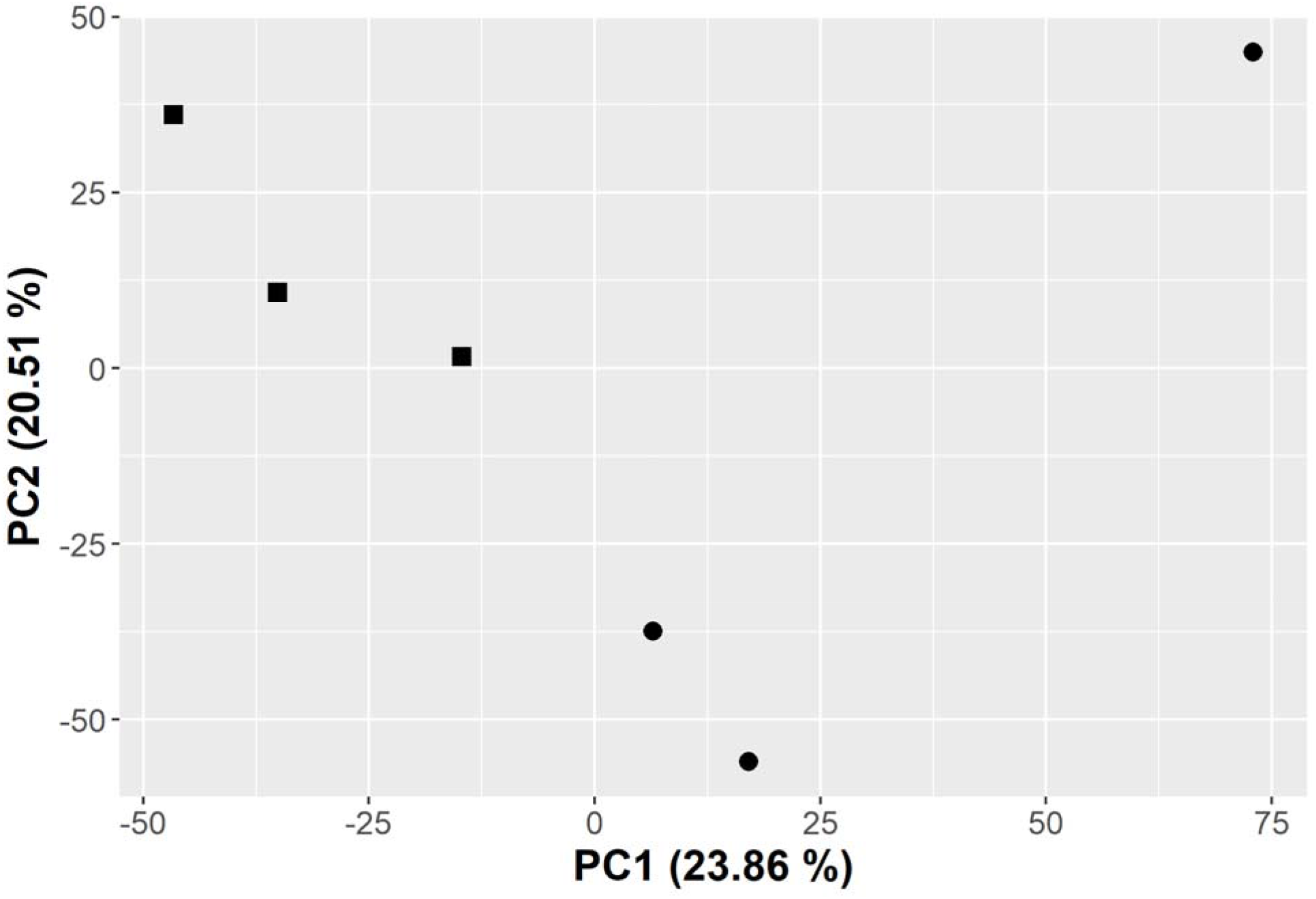
The first two principal components plotted against each other. The percentages in parentheses represent the percent variability explained by the principal component. Circles represent control mice; squares represent mice exposed to hemorrhage followed by cecal ligation and puncture.

A total of ten mice exposed to trauma were part of the survival experiment. A mortality rate of 30% was observed, which is consistent with previous studies using this model. When plotting the first two principal components against each other, the mice who did not survive closely clustered together (Figure 2).

**Figure 2.**
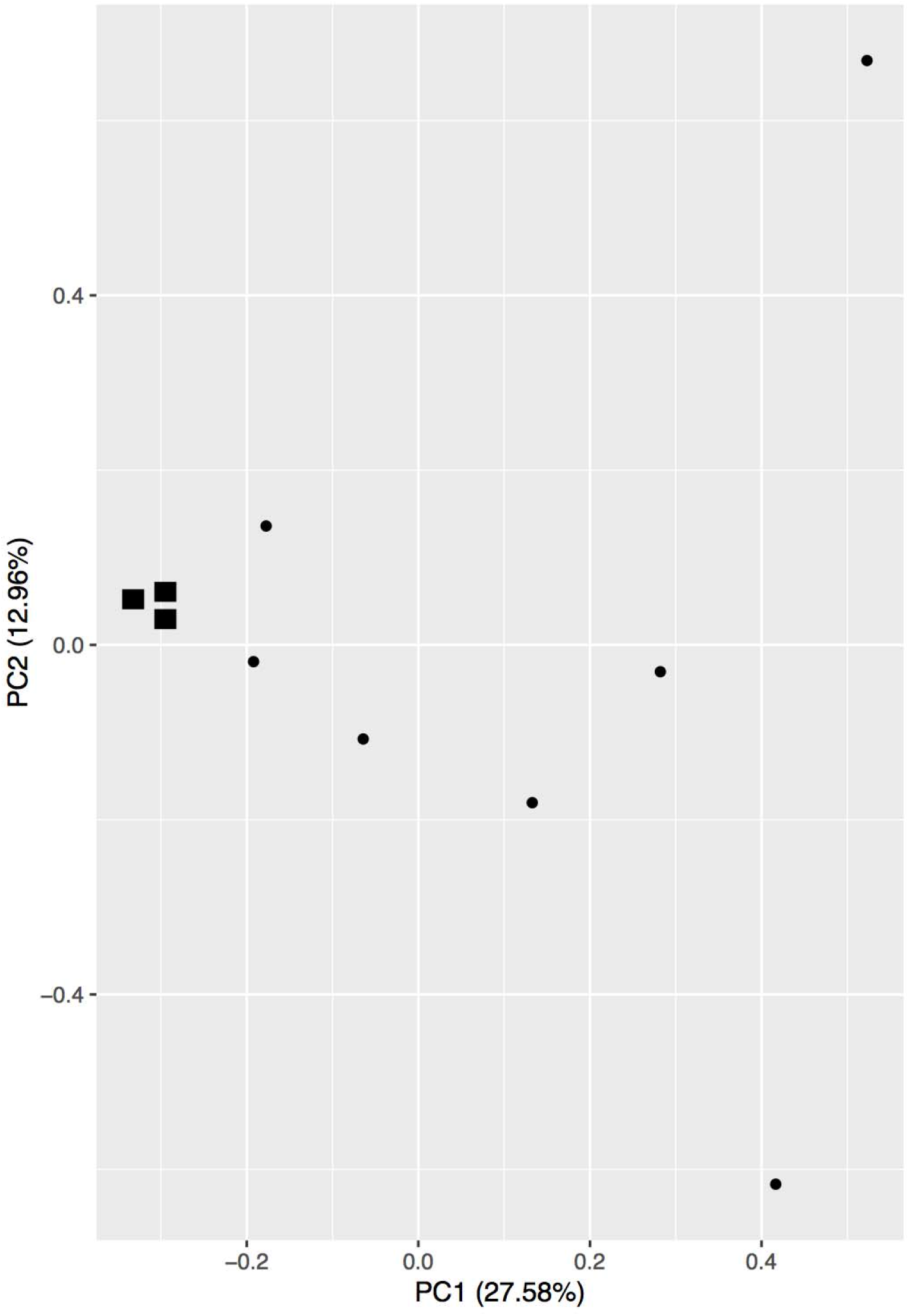
PCA of the survival study. The first two principal components are plotted against each other. The percentages represent the percent variability explained by the principal component. The squares represent mice that died on or before 14 days post CLP, circles represent mice that survived.

## Discussion

In this study, we demonstrated a novel approach of using Deep RNA sequencing to explore differences in the core biological processes that occur in mice exposed to trauma. Most studies using Deep RNA sequencing focus on finding the most significantly different genes or proteins that are aberrantly expressed in a tissue sample. In this study we used a comprehensive measure of transcriptomic entropy to incorporate all pre-mRNA processing events for our analysis. While this approach prevented the identification of a single, tangible set of genes, it enables a birds-eye view on the global dysregulation of alternative pre-mRNA processing in the trauma setting. Alternative pre-mRNA processing entropy has been shown to be different in cancer tissue compared to healthy tissue in a study that used entropy averages of alternative pre-mRNA processing events.^17^ However, to our knowledge, this concept has never been applied to the trauma setting.

Using only transcriptomic entropy values, mice in the trauma group separated from the control group using principal component analysis. This suggests that there is a substantial difference in overall pre-mRNA processing disorder that occurs in the trauma setting compared to healthy controls. This opens the door for further investigation into what drives the difference in pre-mRNA processing entropy, potentially offering new future insights into biological changes that occur in the trauma setting. In addition, this approach is promising for predicting sick versus healthy samples in the trauma setting, offering an additional data point that could be used in future clinical decision making.

While not as clear as in the trauma vs. control experiment, mice that did not survive in the survival study clustered closely together in the principal component analysis (Figure 2). This finding suggests that using RNA splicing entropy has the potential to predict mortality in the trauma setting. In addition, it suggests that differences in alternative splicing may play a role in survival.

The methodology introduced in this study has several strengths as this technique could be applied to a clinical sample to predict disease states or outcomes such as mortality. While the concept of information entropy is not new, its application to alternative pre-mRNA processing, particularly in the trauma setting, is new. By using every single alternative pre-mRNA processing event without manipulation, this method ensures that no information is lost by averaging or other transformations. This is particularly important when examining the level of disorder, where some processing events may see a decrease in entropy while others see an increase. A limitation of this study, is that all the analysis was only done at a single time point. Future work will look at multiple time points to assess if this metric can show transition to recovery. However, this approach does run the risk of incorporating noisy data into the analysis, as no filter is applied to the entropy values. In addition, it is important to note that the changes in entropy seen could be due to changes in the cellular composition of blood in the sick animals versus control. However, this limitation is offset by the sampling of the blood all at the same time after hemorrhage in the survival study analysis. The data presented here is from a single strain of mice who are of similar age; how this metric applies to a heterogeneous human population is not yet confirmed. Finally, the power of this analysis uses all ~300,000 RNA processing events and their entropy values as global measure but we do not know specific events that are responsible for diagnosis of the trauma or prediction of mortality. Future work will attempt to identify causative genes or RNA processing events. The method introduced here is easily transferred to other tissue types or experimental settings. It uses readily available open-source software that is simple to use, with many parts of the analysis that can be conducted on a regular laptop.

## Notes

### Competing Interest Statement

The authors have declared no competing interest.

